# Origin and evolution of papillomavirus (onco)genes and genomes

**DOI:** 10.1101/428912

**Authors:** Anouk Willemsen, Ignacio G. Bravo

## Abstract

Papillomaviruses (PVs) are ancient viruses infecting vertebrates, from fish to mammals. Although the genomes of PVs are small and show conserved synteny, PVs display large genotypic diversity and ample variation in the phenotypic presentation of the infection. Most PVs genomes contain two small early genes *E6* and *E7*. In a bunch of closely related human PVs, the E6 and E7 proteins provide the viruses with oncogenic potential.

The recent discoveries of PVs without *E6* and *E7* in different fish species place a new root on the PV tree, and suggest that the ancestral PV consisted of the minimal PV backbone *E1-E2-L2-L1*.

Bayesian phylogenetic analyses date the most recent common ancestor of the PV backbone to 424 million years ago (Ma). Common ancestry tests on extant *E6* and *E7* genes indicate that they share respectively a common ancestor dating back to at least 184 Ma. In *AlphaPVs* infecting primates, the appearance of the *E5* oncogene 53-58 Ma concurred with i) a significant increase in substitution rate, ii) a basal radiation, and iii) key gain of functions in E6 and E7. This series of events was instrumental to build the extant phenotype of oncogenic human PVs.

Our results assemble the current knowledge on PV diversity and present an ancient evolutionary timeline punctuated by evolutionary innovations in the history of this successful viral family.

## Background

Papillomaviruses (PVs) present a small, circular double-stranded DNA genome with an average size of 8 kbp. A canonical PV genome is organized into three major regions: an upstream regulatory region (URR), an early gene region encoding for three to six proteins (E6, E7, E5, E1, E2, and E4 nested in E2), and a late gene region encoding for two capsid proteins (L2 and L1) (Bravo and Félez-Sánchez, 2015). Proteins in the early region are (among other functions) involved in viral replication and cell transformation, while the capsid proteins self assemble to yield virions and encapsidate the genome. Some of the early genes are dispensable, and the minimal PV genome may contain an URR together with the *E1-E2-L2-L1* genes.

Most PVs are part and parcel of a healthy skin microbiota causing asymptomatic infections in skin and mucosa. In humans, the best studied host, certain human PVs (HPVs) may cause benign lesions, such as skin and genital warts, where the transmission of genital warts occurs primarily via sexual activity (Burchell *et al*., 2006). Only a limited number of evolutionary related HPVs are associated to malignant lesions, that can develop into cancer (Bravo and Alonso, 2004; Schiffman *et al*., 2005). Oncogenic HPVs are a major public health concern as they are responsible for virtually all cases of cervical (99%) and anal cancer (88%), and for a fraction of cancers on the vagina (78%), penis (51%), oropharynx (13-60%, depending on the geographic region) and vulva (15-48%, depending on age) (World Health Organization, 2017). In oncogenic HPVs, the E5, E6 and E7 proteins are directly involved in the onset of cancer. Specific cellular activities of these genes are linked to the virus oncogenic potential and PVs that do not contain these genes are not associated to cancer. In oncogenic PVs, E6 is able to induce degradation of the p53 tumor suppressor protein (Fu *et al*., 2010; Mesplede *et al*., 2012; Sherman *et al*., 1997), while the E7 protein degrades members of the retinoblastoma protein (pRb) family (Dyson *et al*., 1989), which also act as tumor suppressors. The E5 protein in oncogenic PVs is involved in evasion of the immune response and decreases the cellular dependence from external growth factors, thus inducing cell proliferation (DiMaio and Petti, 2013).

PVs were first isolated in mammals, but were later also found to infect birds, turtles, snakes and fish. The first fish PV was recently discovered in skin lesions in a bony fish and named after its host *Sparus aurata papillomavirus 1* (SaPV1) (López-Bueno *et al*., 2016). The SaPV1 genome exhibits an unique organization, as it up to date the only PV that consists of the minimal PV backbone. Moreover, the nucleotide sequences of SaPV1 are so divergent that its discovery led to the recent proposal of reorganising the *Papillomaviridae* taxonomy into two subfamilies: *Firstpapillomavirinae* containing 52 genera and *Secondpapillomavirinae* containing 1 new genus to which SaPV1 belongs.

PVs have evolved in close relationship with their hosts, which allows to perform phylogenetic inference based on host fossil records. However, PV diversity cannot be explained by virus-host codivergence alone. As shown in previous studies (Gottschling *et al*., 2007, 2011b), distantly related PVs infect the same host species, suggesting independent codivergence between viruses within clades and their hosts. The growing number of animal PV sequences available in the online databases allow us to add pieces to the puzzle on the origin and evolution of PV genes and genomes. The modular structure of the PV genome, and the genome organisation of PVs infecting fish reinforce the proposed evolutionary scenario of an ancestral PV that did not contain any of the *E5, E6*, or *E7* genes, which would have been subsequently acquired during PV evolution (García-Vallvé *et al*., 2005).

To better understand how certain HPVs became oncogenic, we have set of a global dating study to look into the evolutionary history of the PV genes and genomes. We have payed special attention to the origin of the PV oncogenes, assessing whether they are monophyletic or have rather originated in several convergent acquisition/loss events. We further look into the roles of these genes and how these novel functions relate to the emergence of oncogenic potential. This paper combining novel data with the previous literature aims to provide insight into the evolution of PVs on a dated scale.

## Materials and Methods

### Data collection and alignments

We collected 354 full length PV genomes (154 animal and 200 human) from the PaVE (pave.niaid.nih.gov, Van Doorslaer *et al*., 2017a) and GenBank (https://www.ncbi.nlm.nih.gov/genbank/) databases (table S1). The previously identified recombinant PVs isolated from Cetaceans (PphPV1-2, TtPV1-7, DdPV1, PsPV1) (Gottschling *et al*., 2011a; Rector *et al*., 2008; Robles-Sikisaka *et al*., 2012) were removed from the data set, leaving us with a data set of 343 full length PV genomes. The *E6, E7, E1, E2, E5, L2* and *L1* genes were extracted and aligned individually at the amino acid level using MAFFT v.7.271 (Katoh and Standley, 2013), corrected manually, and backtranslated to nucleotides using PAL2NAL v.14 (Suyama *et al*., 2006). The alignment was filtered using Gblocks v.0.91b (Castresana, 2000). For tree construction, the *E1, E2, L2* and *L1* were concatenated using a custom perl script. The *E6, E7* and *E5* oncogenes were tested for presence/absence in the data set. The size of these genes was calculated for each PV, and in some cases, the annotation was manually corrected before alignment.

### Phylogenetic analyses and dating

For the concatenated *E1-E2-L2-L1* alignment a ML tree was constructed at the nucleotide level, using RAxML v.8.2.9, under the GTR+Γ4 model, using 12 partitions (three for each gene, corresponding to each codon position) and 1000 bootstrap replicates (fig. S1A). The tree was rooted using SaPV1. Based on the best-known ML tree, 18 calibration points were selected on subtrees where the *E1-E2* an *L2-L1* trees did not show discrepancies and where the host tree matched the PV tree. Calibration point estimates were based on host fossil records from (TimeTree: http://www.timetree.org/). The effect of the calibration points, and therewith forced clades, on the topology of the tree was validated by constructing a ML tree constrained to the calibrations used (fig. S1B) and subsequent comparison to the unconstrained tree using an Shimodaira-Hasegawa test (Shimodaira and Hasegawa, 1999), as implemented in RAxML. The constrained tree (Likelihood: −977756.687001) was not significantly worse than the unconstrained tree (Likelihood: −977683.344298) at the 2% significance level (ΔLH: −73.342703; SD: 32.00218).

Bayesian time inference was performed at the nucleotide level using BEAST v.1.8.3 (Drummond *et al*., 2012), under the GTR+Γ4 model, using twelve partitions, the uncorrelated relaxed clock model (Drummond *et al*., 2006) with a lognormal distribution and a continuous quantile parametrisation (Li and Drummond, 2012), and the Yule Speciation Process tree prior (Gernhard, 2008; Yule, 1925). The constrained ML tree was used as starting tree for time inference. Two independent MCMC chains were run for a maximum of 10^9^ generations and combined when convergence was reached. Chain #1 consisted of 5.249*10^8^ states with an Effective Sample Size (ESS) of 1464, 419, and 4169 for the posterior, prior, and likelihood respectively. Chain #2 consisted of 5.265* 10^8^ states with an ESS of 1227, 360, and 3110 for the posterior, prior, and likelihood respectively. An uncollapsed version of tree figure with the two chains combined can be found in the Supplementary Material (fig. S2).

### Common ancestry test for the *E6* and *E7* oncogenes

To test whether the extant *E6* and *E7* genes have respectively a single common ancestor, we used the software Bali-Phy (Suchard and Redelings, 2006). In this approach, the input data are the unaligned sequences, where the alignment is one of the parameters to be treated as an unknown random variable (de Oliveira Martins and Posada, 2014). We ran our analysis on two different reduced data sets, containing representative species from the different PV clades in fig. 1. Each reduced data set contained 35 sequences representing the different PV crown groups, including the seven PVs infecting Aves and Testudines (grey clade), six Alpha-Omikron PVs (red clade), six Beta-Xi PVs (green clade), six Lambda-Mu PVs (yellow clade), six Delta-Zeta PVs (blue clade), and the four PVs infecting Manatees (black clade). For each data set we ran the analysis for *E6* and *E7*, both separately and concatenated. We assumed the null hypothesis (H0) of Common Ancestry (CA). The Bali-Phy analyses were performed at the amino acid level using the LG substitution model. The likelihood for the CA model was obtained running the software for all the *E6* and/or *E7* sequences together. For the different *Independent Origin* models (IO), we ran the analysis for each group independently. We only considered IO scenarios that were biologically plausible based on the PV tree (fig. 1). For each model, three independent MCMC chains were run for 100,000 iterations. The three runs were combined and checked for convergence. Subsequently, the marginal likelihood was calculated over the three runs using the stabilized harmonic mean estimator. The Bayes Factor for CA is then ΔBF= log[Prob(CA)]-log[Prob(IO)], such that positive values favor CA and negative values indicate IO. As a control, we conducted the same analyses on the *E1* genes, which are monophyletic (table S2).

**Figure 1.**
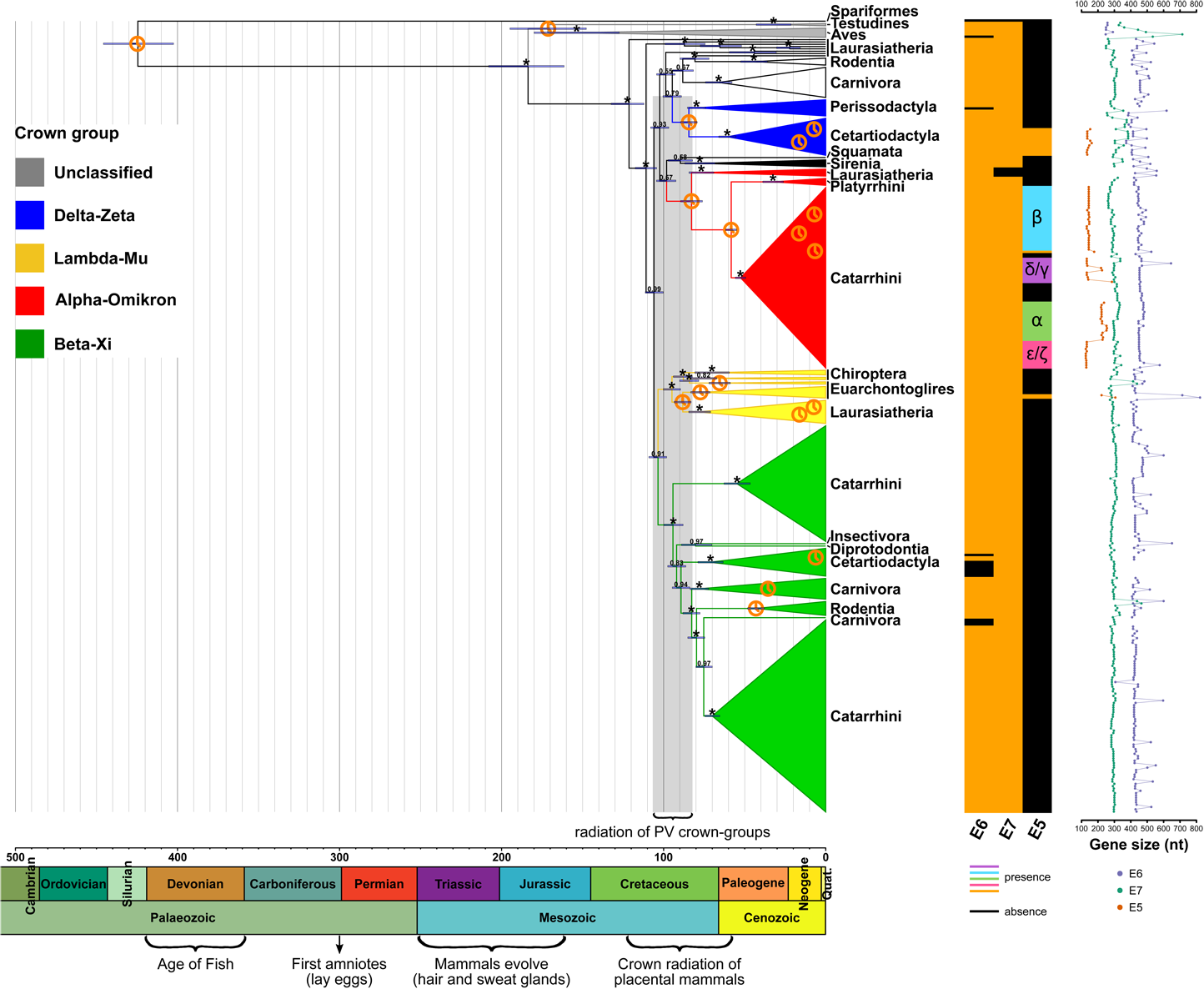
Dated Bayesian phylogenetic tree for a data set containing 343 PVs. The tree was constructed at the nucleotide level based on the concatenated *E1-E2-L2-L1* genes. The scale bar is given in million years ago (Ma). Values at the nodes correspond to posterior probabilities, where asterisks indicate full support. Error bars encompass 95% highest posterior density for the age of the nodes. Clock symbols indicate the nodes used for calibration. Clades are colored according to the PV crown group classification, as indicated in the legend on the left. Next to the tree on the right, the taxonomic host group (superorder, class, order, parvorder, no rank) corresponds to the one in which the clades could best be summarized. Below the tree, a geological time scale is drawn. The matrix next to the taxonomic host groups indicates the presence/absence of the *E6, E7* and *E5* genes for each PV (see legend), and the classification of E5 (*α, β, γ, δ, ε, ζ*) is indicated within the matrix. Next to the matrix, the size of the oncogenes is plotted

To support the results of the Bali-Phy analyses, we performed a random permutation test as described in de Oliveira Martins and Posada, 2016. In this test the sequences for one of the groups are randomly shuffled and statistics are recalculated after realignment with MUSCLE (Edgar, 2004), which tells us how much the original data depart from those with phylogenetic structure partially removed. The statistics used in this test are ML tree lengths and Log Likelihoods, calculated with PhyMLv3.0 (Guindon *et al*., 2010). We reshuffled one of the groups 100 times, each time realigning against the other groups in the data set. For each iteration, the alignment is always optimised and to make the statistics comparable, the same alignment is used for both the IO and CA hypotheses.

### Statistics and graphics

Statistical analyses and graphics were done using R (R Core Team, 2014), with the aid of the packages “ape”, “car”, “ggplot2”, “ggtree”, “stats”, “strap”, “pgirmess”, and “reshape2”. The final display of the graphics was designed using Inkscape v.0.92 (https://inkscape.org/en/).

## Results and Discussion

### Papillomavirus evolution: old ancestry, primary radiation, and secondary diversification

Although the number of animal PV sequences in the databases is growing, the number of available non-mammalian sequences remains low. At the start of this study we recovered five bird PVs, two turtle PVs, one python PV and one fish PV. This fish PV was recovered from lesions in the bony fish gilthead seabream (*Sparus aurata*) (SaPV1) (López-Bueno *et al*., 2016). The genome of SaPV1 is significantly smaller (5,748 kbp) than most previously described PVs (around 8 kbp), and is unique as it is the only PV that contains the minimal PV backbone *E1-E2-L2-L1* while lacking any of the oncogenes (E5, *E6* and E7). This particular genome structure of PVs infecting fish has been confirmed by metagenomic assembly of other fish hosts enriched enriched for circular DNA viruses (Genbank accessions: MH510267, MH616908, MH617143, MH617579), which were made available during the redaction of this manuscript. Besides the unique genome organization of these PVs, the distant sequence relatedness to other PVs suggests a new root on the tree (Spariformes in fig. 1). A recent study dated back this root to 481 Ma, albeit with a broad HPD interval (95% HPD 326 - 656) (Van Doorslaer *et al*., 2017b). In our study, the root of the tree was dated back to 424 Ma (95% HPD: 402 - 446) (fig. 1), which is just before the Devonian period (also often named “Age of Fish”). These results indicate that PVs have an **old ancestry**, so that the ancestral anamniotes were already infected by ancestral PVs in the Palaeozoic, and more specifically in the Silurian period, when bony fish appeared. Millions of years later, in the Mesozoic, we find the last common ancestor of PVs infecting amniotes at around 184 Ma (95% HPD: 161 - 208), followed by a split of PVs infecting birds (Aves) and turtles (Testudines) (grey clade in fig. 1) and PVs infecting mammals (all other PVs with one exception, described below).

The period between the last common ancestor of amniotes and the last common ancestor of mammals corresponds to the evolution of the main traits and skin structures exclusive to mammals, namely hairs, sweat glands, sebaceous glands and milk glands. It has been proposed that the modifications in the proto-mammalian skin environment increased the availability of novel cellular targets for PVs (Bravo and Félez-Sánchez, 2015), so that adaptation to these new niches lead to a **primary radiation** within PVs. A secondary diversification of PVs infecting mammals started around 120 Ma, and the radiation that generated the different PV crown groups dates back to between 106 Ma and 83 Ma, which fits well within the time of crown radiation of placental mammals (fig. 1) (Bininda-Emonds *et al*., 2007). Most PVs infecting mammals have been classified into four different crown groups: Delta-Zeta, Lambda-Mu, Alpha-Omikron, and Beta-Xi (fig. 1). PV crown groups are named after the two most distantly related species therein enclosed (Gottschling *et al*., 2011b). The inferred age for the ancestors of the different PV crown groups is significantly younger than the root age, and significantly different between groups (table 1). This **secondary diversification** event corresponds to the independent co-divergence between viruses and their hosts within each of the major viruses-host clades, generating the enormous diversity of PVs we observe today. Most of these PVs are associated with asymptomatic infections, while some of them cause productive infections, and only a few display carcinogenic potential.

**Table 1.** Inferred node age in million years ago (Ma) for the most recent common ancestors (MRCA) of the different PV clades and for the root of the tree. The rows of the PV crown groups are coloured accordingly (fig. 1), otherwise the taxonomic host group is given. The asterisk indicates the presence of one exception within the clade of PVs infecting mammals, which is a python PV (discussed in the text). The differences between the ancestral node ages of the crown groups as well as the root are significant after performing a Kruskal-Wallis rank sum test (*chi-sq* = 51993, *df* = 5, *p <* 2.2e-16) and a multiple comparison test after Kruskal-Wallis (table S3). Although the inferred times and the posterior distributions for the ancestral Alpha-Omikron and Delta-Zeta as well as the Beta-Xi and Lambda-Mu clades are similar (fig. S3), the significant difference between these groups was confirmed by a Wilcoxon rank sum test (W = 57451000, *p <* 2.2e-16 and *W* = 50158000, *p <* 2.2e-16, respectively).

| PV clade | MRCA age (Ma) | 95% HPD |
| --- | --- | --- |
| root | 424 | 402 – 446 |
| amniotes | 184 | 161 – 208 |
| Aves/Testudines | 171 | 148 – 195 |
| mammals* | 121 | 112 – 132 |
| Lambda-Mu | 95 | 90 – 100 |
| Beta-Xi | 94 | 88 – 100 |
| Delta-Zeta | 84 | 79 – 90 |
| Alpha-Omikron | 83 | 76 – 90 |
| Sirenia | 77 | 68 – 87 |

### Inconsistencies of the current scenario of PVs evolution

The proposed scenario of a biphasic evolution of PVs fits globally well all available phylogenetic analyses (Bravo and Feléz-Sánchez, 2015), including those that we present here. Nevertheless, we observe a number of flaws and anomalies in the PV tree that cannot be explained with our current, linear understanding of PV evolution. Most often, inconsistencies can be imputed to the non-systematic viral sampling, hitherto largely based on economic or leisure interests of the host species, as well as on opportunistic sampling by a reduced circle of veterinarians. Generally, a comprehensive understanding on PV evolution needs to integrate additional evolutionary mechanisms, such as lineage sorting and host switch (Gottschling *et al*., 2011b), which may radically change the relationship between a viral lineage and its host species.

The first anomaly is weak consistency in time inference in the deep nodes of the PV tree, as the inferred age of the MRCA of PVs infecting amniotes (184 Ma, table 1) is more recent than the estimated divergence time between Mammalia and Sauropsida –the two main amniote lines–, which is around 312 Ma (CI: 297 - 326) (estimate derived from 22 studies at http://www.timetree.org/). We did not force any timing interval for this node in our calibrations, so that this result of a “younger-than-expected” node most likely reflects the lack of sufficient phylogenetic information around it, with a long branch connecting it to the root. We will need to increment our sample size for mainly PVs infecting sauropsids and bony fish, to obtain a more accurate time estimate for the MRCA of PVs infecting amniotes.

The second, conspicuous anomaly is the presence of a PV infecting a carpet python; *Morelia spilota papillomavirus 1* (MsPV1) well nested within mammalian PVs (Squamata in fig. 1). The viral genome was retrieved from histologically assessed papilloma-like neoplasias in one python, that tested negative for herpesvirus infections (Gull *et al*., 2012). This PV infecting squamates is not a lonesome exception, as during the writing of this manuscript, the genome of a *Boa constrictor papillomavirus* (BcPV1) was made available (Genbank accession: MH605022). A standard protein BLAST (Altschul *et al*., 1997) identified E1 and L1 of the boa PV as the best hits to the python PV with 58% and 69% of identity, respectively. We have further confirmed that the boa and python PVs are indeed sister taxa in a newly built E1-E2-L2-L1 tree (fig. S4). This clustering supports the phylogenetic position of these two PVs infecting squamates to be close to the unresolved crown of mammalian PV crown groups. The lineages of boas and pythons splitted only 74 Ma (http://www.timetree.org/) and if this timing applies to the corresponding viruses, their MRCA would also fit well in the secondary diversification of PVs. Overall histological assessment of MsPV1 in papillomatous lesions and mostly the finding of BcPV1 strongly suggest that there exists a genuine group of PVs infecting squamates that clusters with mammalian PVs rather than with PVs infecting birds and turtles. We propose that a this lineage emerged after a host switch of an ancestral mammalian PV. Most likely, the evolutionary changes during the secondary PV radiation may have facilitated colonisation of this new ancestral squamate host.

The third anomaly of a solitary PV branching off inconsistently with host phylogeny is a PV genome isolated from a brush-tailed bettong (*Bettongia penicillata papillomavirus 1;* BpPV1), a rare marsupial (Bennett *et al*., 2010). This is the only full-genome report of PVs in marsupials, although fragmentary evidence for PVs infecting marsupials and monotremes has been communicated (Antonsson and McMillan, 2006). We report here BpPV1 (Diprotodontia in fig. 1) to be well-nested within the Beta-Xi crown group, sharing a common ancestor 80 Ma (95% HPD: 70 - 89) with a European hedgehod PV (EePV1; host Insectivora in fig. 1), also solitary in this crown group. Since all other PVs in this crown group have been retrieved from Laurasiatheria, we interpret that a host switch from a PV infecting placental PVs towards marsupials occurred after the emergence of the Beta-Xi crown group, generating this lineage. A systematic screening for PVs in marsupials and monotremes is seriously needed, aiming to populate the viral tree in this important clades of non-placental mammals.

Finally, the fourth anomaly in the PV tree is the poor resolution around the crown of mammalian PVs. Should the null hypothesis of virus-host co-diversification be true, for placental mammals one would expect PVs infecting Xenarthra and Afrotheria to be basal to PVs infecting Euarchontoglires and Laurasiatheria. However, no PV infecting Xenarthrans has been communicated yet, and within Afrotherians only a few PVs infecting manatees (black Sirenia clade in fig. 1) have been reported. These manatee PVs are clearly monophyletic, with an ancestor dating back to 77 Ma (95% HPD: 68 - 87). Although their position within the crown of PVs infecting placental mammals is not well resolved, they are definitely not basal to all PVs infecting Laurasiatherians and Euarchontoglires. Indeed, the MRCA of PVs infecting placental mammals dates back to 121 Ma (95% HPD 112 - 132), and the most basal PVs is consistently a fruit bat PV (Rosettus aegyptiacus PV1, RaPV1) (Rector *et al*., 2006). This virus does not cluster with any other bat PV included in this study, which actually are rather dispersed over the tree (García-Pérez *et al*., 2013, 2014). Once again, we interpret that the lack of appropriate host sampling prevents resolution of the mammalian PV crown.

### The extant *E6* and *E7* oncogenes have respectively a common ancestor, but extant *E5* oncogenes are not monophyletic

We have recently shown that extant *E5* oncogenes do not have a common ancestor (Félez-Sánchez *et al*., 2018). This is not surprising as E5 proteins are only present in few PV clades (fig. 1), and are highly divergent. On the contrary, both the *E6* and *E7* oncogenes are present in most PV genomes, and are less divergent than *E5* (Bravo and Alonso, 2004). Interestingly, some PVs are lacking either *E6* or *E7* (presence/absence matrix in fig. 1). A bunch of PVs lacking *E7* (MrPV1, PphPV4, SsPV1, UmPV1) infect Laurasiatherian hosts and belong to the the Alpha-Omikron PV crown group. All other members of this crown group infect Primates and present both *E6* and *E7*. On the other hand, PVs lacking *E6* are more disperse along the tree and belong into different crown groups: one parrot (PePV1) PV in the grey clade, one donkey PV (EaPV1) in the Delta-Zeta crown group, and eight bovine PVs (BPV3, BPV4, BPV6, BPV9, BPV10, BPV11, BPV12, BPV15) and three human PVs (HPV101, HPV103, HPV108) in the Beta-Xi crown group.

Regarding the size of the oncogenes (indicated next to the tree in fig. 1), *E6* is small (median: 253.5 nt; Q_1_: 249.0 - Q_3_: 258.0) in the genomes of PVs infecting birds and turtles (grey clade), while it is of a double size (median: 438.0 nt; Q_1_: 420.0 - Q_3_: 462.0) in the genomes of all PVs infecting mammals. This increase in size correlates with the presence of a second *E6* zinc binding motif domain (Zanier *et al*., 2012), which could have appeared after duplication of the first original motif and transformed the original homodimer into an internal dimer (Suarez and Trave, 2018).

As described above, during the production of this manuscript four novel fish PV genomes became available at the Genbank database, infecting 3 new bony fish species (1 rainbow trout, 1 red snapper and 2 haddocks). None of the genomes of these novel fish PVs contain any of the oncogenes. For the now five fish PVs, the most conserved *E1* and *L1* genes present a high sequence diversity. To see whether the novel fish PVs cluster together with the ancestral SaPV1, we recalculated our ML tree based on the concatenated *E1-E2-L2-L1* sequences. Indeed, we found that the all the fish PVs are monophyletic fig. S4).

The pattern of presence/absence of *E6* and *E7* in extant PVs demands an evolutionary explanation. One hypothesis proposes that the MRCA of all PVs already contained these ORFs and invokes six independent repeated loss events for *E6* in different PV lineages (Van Doorslaer and McBride, 2016), including in the lineage leading to extant fish PVs, and one gene loss event for *E7* in another PV lineage. An alternative hypothesis, more parsimonious with the absence of *E6* and *E7* in fish PVs, proposes an ancestral PV genome spanning only the minimal arrangement *E1-E2-L2-L1*, the gain of the ancestral *E6* and *E7* genes in the lineage of amniote PVs, at least 184 Ma, followed by five independent loss events for *E6* and one loss event for *E7*.

To unravel a piece of the evolutionary history of *E6* and *E7*, we performed common ancestry tests as described in de Oliveira Martins and Posada, 2014 using the program Bali-Phy (Suchard and Redelings, 2006). The advantage of this approach is that the alignment and the phylogeny are estimated at the same time, reducing the bias towards obtaining Common Ancestry (CA) introduced by the alignment step. We ran our analyses on two reduced data sets containing representatives from each PV clade. We tested different hypotheses supporting either CA or independent origin (IO) for *E6* and *E7* separately as well as concatenated. We only considered IO scenarios that were biologically plausible based on the PV tree, leading us to eight different hypotheses (H0-H7), as displayed in tables 2 and 3. We assumed the null hypothesis (H0) of common ancestry (Materials and Methods). For the independent origin scenarios (H1-H7) we performed the analysis separately for each group, where the sum of the marginal likelihood of these groups represents that of the hypothesis tested. As an example, for H2, we ran one analysis for the Aves/Testudines clade (grey), one analysis for the Delta-Zeta crown group (blue clade), and one analysis for the Lambda-Mu crown group (yellow clade), Alpha-Omikron crown group (red clade), the Sirenia clade (black), and the Beta-Xi crown group (green clade) together. Then we calculated ΔBF= log[Prob(CA)]-log[Prob(IO)], such that positive values favor CA and negative values indicate IO. The overall results suggest (table 2 and 3) that extant *E6* and *E7* share respectively a common ancestor, independently of whether the analyses were performed on the concatenated *E6-E7*, or on *E6* and *E7* alone. Although convergence was reached between the three independent MCMC runs for all groups within the hypotheses tested, we perform an additional permutation test for stronger evidence of CA. This test was performed as described in de Oliveira Martins and Posada, 2016, where the columns of the alignment for one of the groups are randomly shuffled, such that the alignment is preserved within the group but disrupted between groups. After shuffling the alignment is always optimised for the original data set, so that the ML tree can be estimated and summary statistics can be calculated (see Materials and Methods). The results of the permutation test support the hypothesis of CA as the most-likely scenario (file S1. Nevertheless, this test also reveals that the second-best supported model −H1: IO for *E6* and *E7* in the Aves/Testudines clade– is not significantly worse. Thus, although the best scenario is common origin for all extant *E6* and *E7* genes, we cannot reject the hypothesis that *E6* and *E7* in extant PVs infecting birds and turtles have originated independently from the *E6* and *E7* genes in extant PVs infecting mammals. Only a denser sampling of PV genomes from different large amniote clades outside mammals, as well as from monotremes and marsupials may allow distinguish between these conflicting hypotheses.

**Table 2.** Testing for common ancestry of E6 and E7 on Reduced Data set 1.

| Model | E6E7 |  | E6 |  | E7 |  |
| --- | --- | --- | --- | --- | --- | --- |
| | P(data\|M) | $\Delta BF$ | P(data\|M) | $\Delta BF$ | P(data\|M) | $\Delta BF$ |
| H0: (grey-blue-yellow-red-black-green) | -19083.055 | 0 | -10960.151 | 0 | -8239.297 | 0 |
| H1: grey+(blue-yellow-red-black-green) | -19425.259 | 342.204 | -11008.514 | 48.363 | -8300.605 | 61.308 |
| H2: grey+blue+(yellow-red-black-green) | -19595.075 | 512.020 | -11102.586 | 142.435 | -8391.373 | 152.076 |
| H3: grey+blue+yellow+(red-black-green) | -19883.098 | 800.043 | -11266.682 | 306.531 | -8515.244 | 275.947 |
| H4: grey+blue+yellow+red+(black-green) | -20108.285 | 1025.230 | -11390.535 | 430.384 | -8644.525 | 405.228 |
| H5: grey+blue+(red-black)+(green-yellow) | -19850.013 | 766.958 | -11239.646 | 279.495 | -8541.050 | 301.753 |
| H6: grey+blue+red+black+(green-yellow) | -22355.489 | 3272.434 | -11352.869 | 392.718 | -8638.945 | 399.648 |
| H7: grey+blue+yellow+red+black+green | -20317.714 | 1234.659 | -11503.409 | 543.258 | -8748.908 | 509.611 |

**Table 3.** Testing for common ancestry of E6 and E7 on Reduced Data set 2.

| Model | E6E7 |  | E6 |  | E7 |  |
| --- | --- | --- | --- | --- | --- | --- |
| | P(data\|M) | $\Delta BF$ | P(data\|M) | $\Delta BF$ | P(data\|M) | $\Delta BF$ |
| H0: (grey-blue-yellow-red-black-green) | -19083.055 | 0 | -10847.336 | 0 | -8106.947 | 0 |
| H1: grey+(blue-yellow-red-black-green) | -19165.438 | 82.383 | -10915.029 | 67.693 | -8153.371 | 46.424 |
| H2: grey+blue+(yellow-red-black-green) | -19406.063 | 323.008 | -11020.832 | 173.496 | -8281.371 | 174.424 |
| H3: grey+blue+yellow+(red-black-green) | -19676.573 | 593.518 | -11174.707 | 327.371 | -8424.371 | 317.424 |
| H4: grey+blue+yellow+red+(black-green) | -19921.623 | 838.568 | -11306.744 | 459.408 | -8539.261 | 432.314 |
| H5: grey+blue+(red-black)+(green-yellow) | -19682.881 | 599.826 | -13601.982 | 2754.646 | -8411.536 | 304.589 |
| H6: grey+blue+red+black+(green-yellow) | -19886.211 | 803.156 | -13704.295 | 2856.959 | -8511.299 | 404.352 |
| H7: grey+blue+yellow+red+black+green | -20121.000 | 1037.945 | -11412.956 | 565.620 | -8642.184 | 535.237 |

### The emergence of an oncogenic potential in certain human PVs

A well-substantiated body of scientific literature suggests that the oncogenic potential of certain human PVs lies on the perturbation of key cellular checkpoints by the E6 and E7 proteins. However, although most PVs contain these genes, only a few of them are actually associated to cancer. Among the more than 220 HPVs, around 20 closely related *AlphaPVs* have been classified by the International Agency for the Research on Cancer (IARC) as carcinogenic or potentially carcinogenic to humans. Apart humans, most other cancer cases associated to PVs are rare cases: EcPV2 in penile cancers in stallions (Scase *et al*., 2010), RaPV1 in basosquamous carcinoma in the Egyptian fruit bat (Rector *et al*., 2006), or RrPV1 in a nasolabial tumour in a free-raning chamois (Mengual-Chuliá *et al*., 2014). A more specific rare case is represented by bovine PVs (BPVs). These PVs induce benign tumours of cutaneous or mucosal epithelia in the cattle, however in the case of BPV1 (and less common BPV2), a hosts switch reveals its oncogenic potential, as in horses BPV1 can give rise to malign fibroblastic tumours (sarcoids) (Nasir and Campo, 2008). The later classified BPV13 has also been found in equine sarcoids (Lunardi *et al*., 2013). Nevertheless, BPV1 is as well detected in the skin and blood of healthy horses (Bogaert *et al*., 2005; Federica *et al*., 2015), and suggests that equine-adapted BPV1 strains with the occurrence of viral variants exist (Federica *et al*., 2015; Trewby *et al*., 2014). It has been proposed that BPV1 variant sequences are associated to either a benign or malign phenotype by altering the expression of the E5 protein (Chambers *et al*., 2003). Besides this host switch from a bovine PV to an equine host, eighth different equine PVs have been described (PaVE: pave.niaid.nih.gov, Van Doorslaer *et al*., 2017a). *Equine papillomavirus 2* (*EcPV2*) has been detected in genital squamous cell carcinomas (SCCs) and healthy genital mucosa (Bogaert *et al*., 2012; Sykora *et al*., 2017). However, odd ratios for the presence of viral material in diseased vs healthy animals suggests that that EcPV2 contributes indeed to the onset and progression of genital SCCs in horses (Sykora and Brandt, 2017). Preliminary findings further suggest that EcPV2 resides in infected cells as virions, viral episomes and integrated viral DNA (Sykora *et al*., 2017), similar to cancer associated HPVs in humans (Doorbar *et al*., 2012). This seems to be an example of convergent evolution, where EcPV2 and oncogenic human PVs both evolved independently analogous mechanisms to stimulate the development of PV-associated cancers.

Within the clade of PVs infecting primates (*AlphaPVs*: Catarrhini in red clade in fig. 1), the E5 proteins are classified into different groups (E5*α*, E5*β*, E5*γ*, E5*δ*, E5*ε*, E5*ζ*) according to their hydrophobic profiles and phylogeny (Bravo and Alonso, 2004). The *AlphaPVs* divide in three subclades with three different clinical presentations: cutaneous warts, genital warts and mucosal lesions (fig. 2). The presence of a given E5 type strongly correlates with the clinical presentation of the corresponding PV infection: E5*α* is associated to malignant mucosal lesions, E5*β* is associated to benign cutaneous lesions, and the two putative proteins E5*γ* and E5*δ* are associated to benign mucosal lesions (Bravo and Alonso, 2004). The appearance of the *E5* proto-oncogene in the ancestral *AlphaPV* genome can be dated back to between 53 and 58 Ma (fig. 2), and concurred with an event that was instrumental for the differential oncogenic potential of present-day human PVs. One hypothesis is that the appearance of *E5* triggered an adaptive radiation that generated the three viral lineages with different clinical manifestations (Bravo and Félez-Sánchez, 2015). Nevertheless, we have recently shown that not all E5s have a common ancestor (Félez-Sánchez *et al*., 2018). We interpret that E5s evolved *de novo* out of an originally non-coding region that integrated between the early and the late genes in the genome of an ancestral *AlphaPV* lineage. Interestingly, at the time of the integration of this non-coding region, we observe an acceleration of the evolutionary rate in the corresponding branch, two times higher than the overall PV substitution rate (fig. 2). These results suggest that the accommodation of the E5s in this region and the promotion of an adaptive radiation, where certain E6 (and probably also E7) proteins acquired the ability to degrade tumor suppressor proteins and facilitate the development of cancer in different tissues.

**Figure 2.**
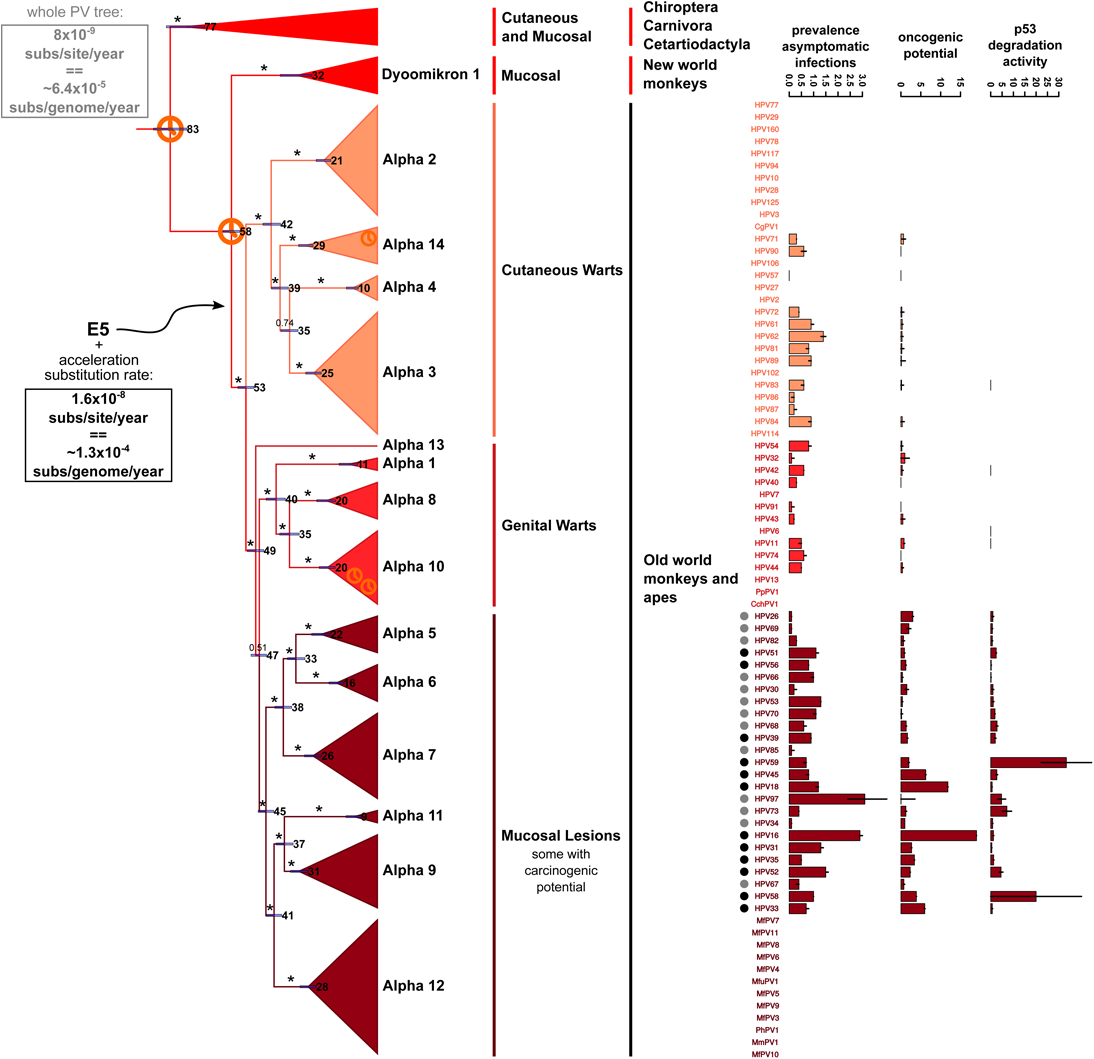
This tree is a zoom of the Alpha-Omikron PV crown-group in fig. 1 above. Values at the nodes correspond to posterior probabilities, where asterisks indicate full support. Error bars encompass 95% highest posterior density for the age of the nodes, next to the error bars the median node age is given, in millions of years ago (Ma). Clock symbols indicate the nodes used for calibration. A black arrow indicates the timing for the emergence of *E5* gene in the ancestral PV genome, between 53 and 58 Ma. Boxes display the average evolutionary rate for the complete PV tree (in grey) or for the *AlphaPV* subtree after the emergence of *E5* (in black). On the right side of tree, the different PV species, the clinical presentation, and host taxonomy are given. Dots label HPVs that have been classified by the IARC as carcinogenic to humans (black dots, group I) or probably/possibly carcinogenic to humans (grey dots, groups IIa and IIb). The three barplots on the right represent: a) the worldwide prevalence of each HPV in women with normal cervical cytology with error bars indicating the 95% confidence interval; b) the oncogenic potential for each HPV, proxyed as the ratio between the prevalence of each HPV in cervical cancers divided by the prevalence in normal cervical cytology) with error bars indicating the 95% confidence interval; c) the E6-mediated p53 degradation activity, expressed as the inverse value of the EC50 in ng of E6 protein needed to degrade cellular p53, with higher values indicating an enhanced potential of E6 to degrade p53; error bars indicate an approximate of the standard error of the mean. The first two columns of the barplots contain data obtained from the ICO/IARC HPV Information Centre (http://www.hpvcentre.net/), while the third column contains data obtained from Mesplede *et al*., 2012. For the raw data of the barplots see table S4.

The oncogenic potential of human PVs strongly matches viral phylogeny Bravo and Alonso, 2004; Schiffman *et al*., 2005. The potential of p53 degradation by E6 proteins from *AlphaPVs* is highly correlated with such phylogenetic grouping (Fu *et al*., 2010), which has suggested a mechanistic basis for the connection between phylogeny and oncogenicity. Although E6-mediated p53 degradation has always been considered one of the hallmarks of HPV-mediated cervical cancer (IARC, 2012), the connection between molecular mechanism and infection phenotye remains unclear. First, E6 proteins from non-oncogenic HPVs, notably HPV71, can also induce p53 degradation (Fu *et al*., 2010). Second, rare cases of malignancy associated to non-oncogenic HPVs, whose E6 proteins do not degrade p53, are well documented (Guimerà *et al*., 2013). Finally, third, when the E6-mediated p53 degradation activity has been finely quantified Mesplede *et al*., 2012, a more complex picture was revealed. Our thought-provoking results displayed in fig. 2 clearly show that there is no correlation between the oncogenic potential for HPVs in cervical cancer, and the E6-mediated p53 degradation activity (Pearson’s product-moment correlation = −0.0303016, *t* = −0.14852, *df* = 24, *p* = 0.8832). Cogent examples are HPV16 and HPV18, which display the highest oncogenic potential (proxyed as the ratio between worldwide prevalence in cervical cancers and worldwide prevalence in women with normal cervical cytology,(data obtained from the ICO/IARC HPV Information Centre), but whose corresponding E6 proteins are not specially efficient at inducing p53 degradation (proxyed as the inverse of the EC50 concentration of E6 needed to degrade 50% of the cellular p53 protein, data obtained from Mesplede *et al*., 2012). For comparison, E6 from HPV58 requires 17-times less concentration to achieve the same p53 degradation effect as E6 from the closely related HPV16, and E6 from HPV59 requires 53-times less concentration to achieve as much p53 degradation as E6 from the closely related HPV18. Other factors, such as mRNA splicing and the presence of particular spliced E6 isoforms specific to oncogenic HPVs (Schmitt *et al*., 2010), may play essential roles in defining the overall oncogenic potential of the different HPVs.

## Conclusion

In this study we revisited the evolution of PVs using phylogenetic dating on the largest and most diverse data set compiled to date. The evolutionary scenario with the best explanatory power proposes an old ancestry for PVs, a primary radiation event that led to the generation of the different main lineages, a second radiation of the different lineages together with the expansion of their hosts, and a third radiation event specific to *AlphaPVs* after the emergence of the *E5* proto-oncogene. We identify further a number of anomalies in the PV tree that are inconsistent with this overall scenario. Some of these inconsistencies can be explained by lineage sorting and host switch events. We also show for the first time that the *E6* and *E7* oncogenes may have a common ancestor, although the alternative hypothesis of *E6* and *E7* from mammalian PVs and from Aves/Testudines PVs cannot be rejected due to the still scarce sampling. Overall, resolution of the deep nodes and fine support for the main scenario will require systematic sampling of PVs from other anamniotes, from amniotes other than mammals, from mammals other than placental mammals, and from placental mammals other than Laurasiatherians and Primates. We need to humbly admit that we are still far from understanding why PV-induced cervical cancers seem to be restricted to humans, as well as from identifying the molecular differences between closely related viruses underlying the enormous variance in the epidemiology of oncogenic PVs.

## Supplementary Material

Supplementary tables S1-S4, figures S1-S4, and file S1 are available online. The data from the Bali-Phy analyses and the random permutation test are available at http://gitlab-bioinfo.ird.ir/awillemsen/ONCOGENEVOL_E6E7.

## Authors’ contributions

AW designed the study, carried out the data collection and analyses, and drafted the manuscript; IGB designed the study, coordinated the study and helped draft the manuscript. All authors gave final approval for publication.

## Acknowledgements

We are grateful to the genotoul bioinformatics platform Toulouse Midi-Pyrenees (Bioinfo Genotoul) and the IRD bioinformatics platform for providing computing and storage resources.

## Abbreviations

CA: : Common Ancestry
IARC: : International Agency for the Research on Cancer
ICTV: : International Committee on Taxonomy of Viruses
IO: : Independent Origin
ESS: : Effective Sample Size
HPD: : Highest Posterior Density
Ma: : Million years ago
MRCA: : Most Recent Common Ancestor
PV: : Papillomavirus
SCC: : Squamos Cell Carcinoma
SEM: : Standard Error of the Mean

